# The role of division stochasticity on the robustness of bacterial size dynamics

**DOI:** 10.1101/2022.07.27.501776

**Authors:** César Nieto, Juan Carlos Arias-Castro, Carlos Sánchez, César Vargas-García, Abhyudai Singh, Juan Manuel Pedraza

## Abstract

Variables of bacterial division such as size at birth, growth rate, division time, and the position of the septal ring, all vary from cell to cell. Currently, it is unknown how these random fluctuations can combine to produce a robust mechanism of homeostasis. To address this question, we studied the dynamics of the cell division process from both experimental and theoretical perspectives. Our model predicts robustness in division times as sustained oscillations in metrics of the cell size distribution, such as the mean, variability, and the cell size autocorrelation function. These oscillations do not get damped, even considering stochasticity in division timing and the cell size at the beginning of the experiment. Damping appears just after inducing stochasticity in either the septum position or the growth rate. We compare the predictions of the full model with the size dynamics of *E. coli* bacteria growing in minimal media using either glucose or glycerol as carbon sources. We observe that growth in poorer media increases the noise in both partitioning position and growth rate. This additional noise results in oscillations with more damping. Although intracellular noise is known as a source of phenotypic variation, our results show that it can play a similar but subtler role in maintaining population-level homeostasis by causing rapid desynchronization of cell cycles..

## Introduction

To achieve proper function, cells must maintain stability in their physiological variables. To maintain this stability, also known as homeostasis, they must synchronize all self-regulating processes [1–4]. Recently, great effort has been made to understand cell size homeostasis [5–7]. However, some questions remain open. For example, it is unclear how the molecular mechanisms that maintain size homeostasis work together despite showing random fluctuations [8].

Recent studies predict that precise control of growth and division processes in exponentially growing cells can lead to oscillations in cell size autocorrelation function (ACF) [9]. These oscillations can be related to a synchronization of the cell cycle for cells that descend from a common ancestor [10]. This synchronization could, in turn, affect size-dependent variables such as protein concentration [11, 12]. Thus, precise size control comes with a dilemma: In terms of maintaining homeostasis, it would be best for the cell to have a growth and division cycle as reliable as possible, but if this cycle were deterministic (perfect control), the cells in a clonal population would be correlated at any longer times. This would mean that the growth medium itself would have periodic fluctuations in concentrations occurring at relevant timescales for metabolic control as the synchronized population grows and divides.

Of course, this is not what is observed experimentally: cell growth and division are inherently stochastic processes [11, 13–15]. This randomness leads to rapid desynchronization of cell cycles [9]. To determine exactly how the different sources of stochasticity affect this desynchronization, we developed a stochastic model based on a continuous division rate [9, 15–19] and describe the stochastic process of cell growth and division that incorporates different sources of noise. At the same time, we have experimentally observed the size dynamics of hundreds of *E. coli* bacteria in Mother Machine microfluidic devices [20] using minimal media with different carbon sources. We observed damped oscillations with the amount of damping depending on the richness of the medium.

In the first part of this article, we present a theory that predicts the oscillations in the cell size ACF. We will also explain how to add some biologically essential variables. These variables can be the division strategy [16, 21] and a number of cycle stages to trigger division [8, 15]. And we show how other sources of noise, such as fluctuations in the growth rate [14] or randomness in the position of the septum ring [22, 23] can damp these oscillations. We then present observations of this damping in the experiments mentioned earlier. We end with a discussion of some consequences of this phenomenon.

## Modeling size dynamics with stochasticity in division timing

In this section, we revisit a known model that describes division as a time-continuous stochastic process [9]. Later, it will be necessary to add the relevant sources of noise.

### Growth and division as continuous-time processes

As presented in Figure 1A, we asume that a cell has size *s* that elongates exponentially over time *t* :

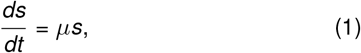

with a growth rate *µ* (or equivalently, with a doubling time *τ* = ln(2)*/µ*).

**Figure 1:**
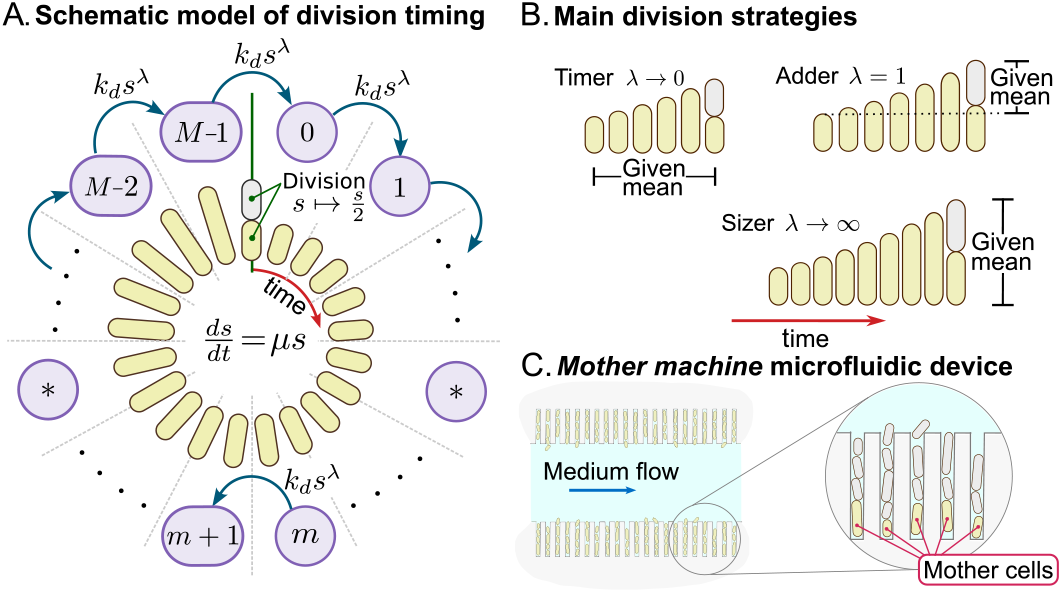
Schematic model, the main strategies of division and Mother machine. **A**. The division is modeled as a stochastic process that occurs after the completion of *M* stages. During division, the cell resets the stages to zero. The transition rate between any stage *m* to *m* + 1 is *k*_*d*_ *s*^*λ*^ where *k*_*d*_ is a constant, *s* is the cell size, and *λ* > 0 is an exponent. During the stages, the cell grows exponentially with a growth rate *µ*. During division, the cell size *s* is halved. **B**. Depending on *λ*, different division rules can be defined. *λ* →0 describes the *timer* where cells, on average, wait a given mean time to divide. *λ* = 1 describes the *adder* where cells add, on average, a given size to divide. *λ* → ∞ describes the *sizer* where cells, on average, divide at a given mean size. **C**. In the *Mother machine* microfluidic device, bacteria are trapped at the bottom of a dead end channel (mother cells). Their descendants are discarded (gray).

As we present in Figure 2A, after each division, one of the two descendants cells is randomly discarded (gray) and the other (yellow) is continuously tracked. This is what we observe in our experiments using the *Mother Machine* microfluidic device (see Figure 1C) [17, 18, 24].

**Figure 2:**
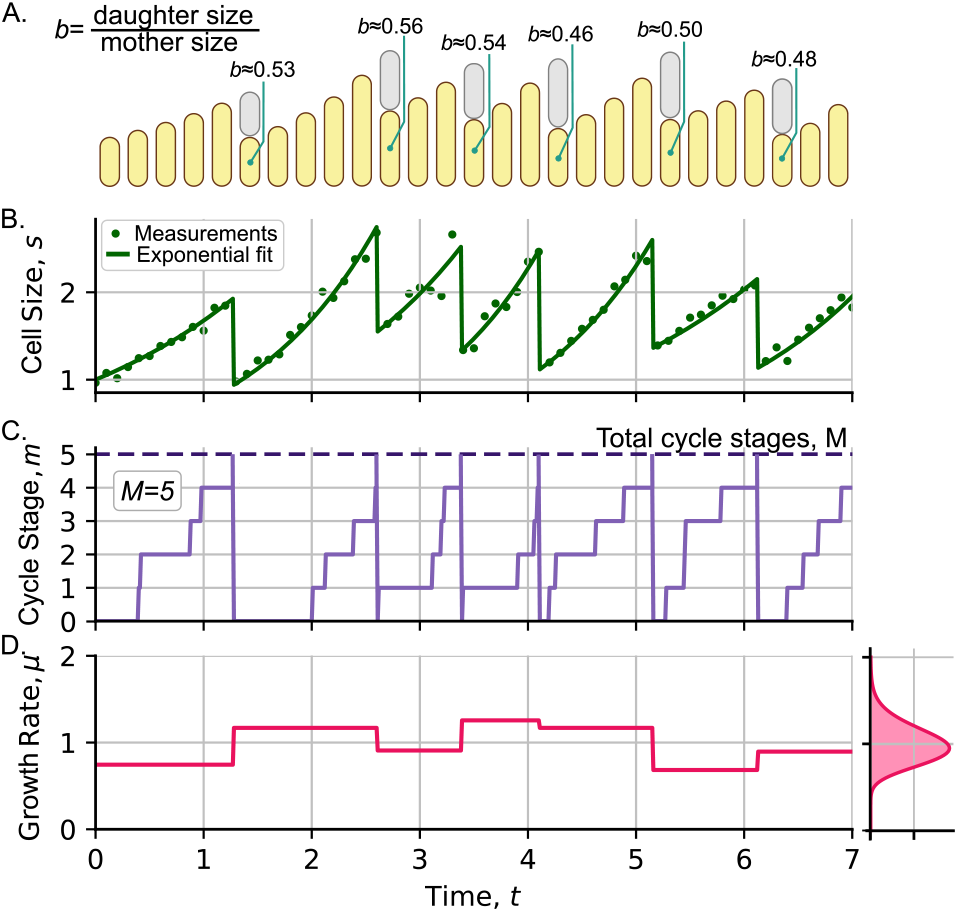
Main variables used for describing the cell size dynamics. **A**. Schematic kymograph of cell size over time. After each division, one of the descendants cells is discarded (gray), while the other is tracked (yellow). The partitioning ratio *b*, the ratio between the size of the daughter cell and its mother just after division, is a random variable i.i.d. chosen from a beta distribution with a given coefficient of variation *CV*^2^(*b*). We also show *b* for some cycles. **B**. Cell size measurements (dots) can be fitted to an exponential function of time (line). **C**. Division is triggered when the cell completes a fixed quantity *M* of cycle stages (in this case *M* = 5, dashed line). **D**. During each cycle, we assume that the growth rate *µ* is constant and is a random variable i.i.d. chosen from a gamma distribution (left) with a given coefficient of variation *CV*^2^(*µ*).

If the size is perfectly halved during division, the cell size *s*(*n*; *t*) after *n* ∈ ℕ divisions at time *t* is [25]:

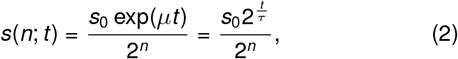

where *s*_0_ is the initial cell size, that is, the cell size at *t* = 0.

Once the growth conditions are defined, the next process to describe the size dynamics is division regulation. Recent models suggest that division is triggered after the occurrence of *M* ∈ *N* intermediate steps that represent the cycle stages [8, 15, 26]. Let *m* ∈ {0, 1, …, *M*} be the current cycle stage of the cell (see Figure 1A and Figure 2C). The transition rate *h*(*s*) from state *m* to *m* + 1 can be modeled as proportional to a power of *s* [15, 18, 19, 27]:

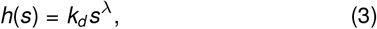

where *k*_*d*_ is a constant and *s* satisfies (2). *λ* > 0 is a parameter that defines the division strategy, which is a concept that we will explain later.

Let *P*_*m,n*_(*t*) be the probability that the cell completes *m* cycle stages and *n* divisions up to time *t*. This probability can be estimated using the rate (3) using a master equation [9]:

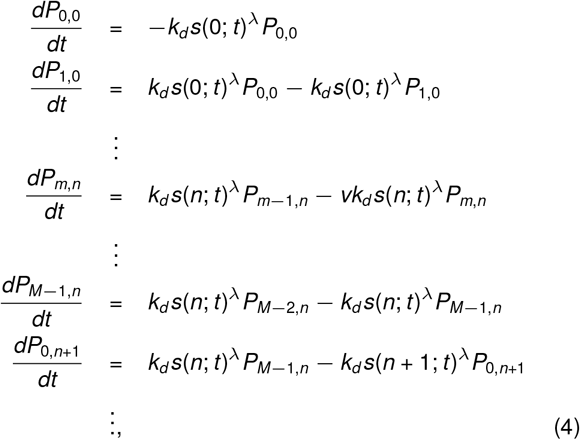

where *P*_*M,n*_ := *P*_0,*n*+1_ (see Figure 1A). This is given since during a division (when *m* reaches *M*), *m* is reset to zero and *n* increases by 1.

### The division strategy

Consider that the cell starts exactly after one division, that is, at *m* = 0 and the division counter is set to *n* = 0. *s*_0_ can take any arbitrary value *s*_*b*_, also known as the size at birth. Following the methods explained in previous studies [9], the probability density *ρ*(*s*_*d*_|*s*_*b*_) of the size at division *s*_*d*_, given the size at birth *s*_*b*_ in a cell cycle, satisfies:

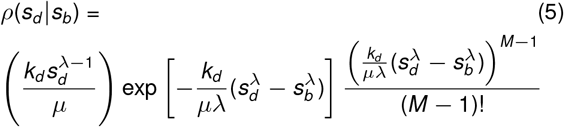

Defining the added size before division Δ = *s*_*d*_ *− s*_*b*_, we observe that if *λ* = 1, (5) reduces to a gamma distribution for the variable Δ. The mean value ⟨Δ⟩ is independent of the size at birth *s*_*b*_, but is related to *µ, M*, and *k*_*d*_ by the expression [18]:

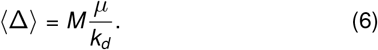

This property, where the mean added size ⟨Δ⟩ does not depend on the size at birth *s*_*b*_ defines the *adder* division strategy [17, 21, 24]. Other strategies (see Figure 1B), such as the *sizer* (a slope of -1 in Δ vs. *s*_*b*_) can be obtained at the limit *λ* → ∞. The *timer* strategy (a slope of +1 in Δ vs. *s*_*b*_) is obtained in the limit *λ* → 0, while intermediate strategies such as the *sizer-like* (a slope between -1 and 0), observed in slow-growing bacteria [21, 22, 28] are obtained with 1 *< λ <* ∞, and *timer-like* strategies (a slope between 0 and 1), found in bacteria such as *C. crescentus* [29], are obtained when 0 *< λ <* 1. It is worth mentioning that there are other possible explanations for the origin of sizer-like strategies [8, 28, 30] that cannot be easily coupled with this theoretical framework.

Given the division strategy, we define 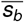 roughly as the average size at birth when the division process reaches stability (technical details are discussed in [9, 18]). After normalizing all cell sizes by 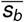, we can compare this theory with experiments.

### Oscillations in central moments of size distribution after synchronization

To calculate the probability density *ρ*(*s*|*t*) of having cell size in the interval (*s, s* + *ds*) at time *t*, for simplicity, we assume that all cells started at *t* = 0 with size *s*_0_. This corresponds to the PDF of the initial cell size:

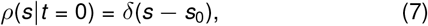

with *δ*(*X*) being the Dirac delta distribution.

The probability *P*_*n*_(*t*) that the cell has divided *n* times up to time *t* is obtained by the marginal sum:

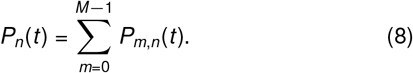

Using (7) and (4), *ρ*(*s*|*t*) is given by the following:

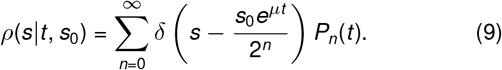

PDF (9) corresponds to the sum of the weighted Dirac delta distributions *δ*(*X*) with positions centered on the sizes (2). The expected value ⟨*s*⟩ of the cell size and its variance var(s) follow:

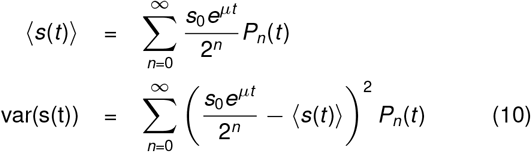

Figure 3 shows some examples of the mean size trajectories ⟨*s*⟩ and the size variability *CV*^2^(*s*) = var(*s*)*/* ⟨*s*⟩ ^2^ for different total stages *M* and division strategies *λ* in (3) under the initial condition *P*_*m,n*_(*t* = 0) = *δ*_*M*,0_*δ*_*n*,0_ with *δ*_*i,j*_ being the Kronecker delta.

**Figure 3:**
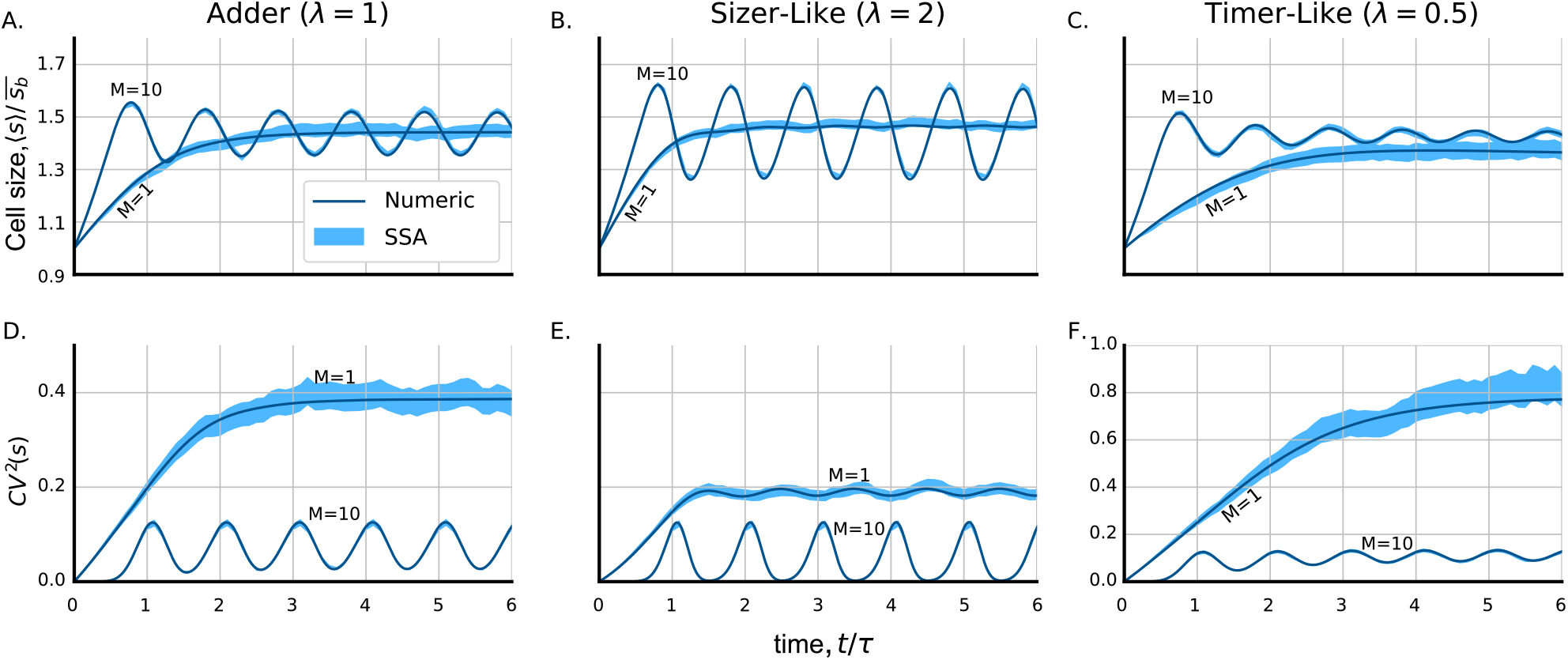
Theoretical dynamics of the size distribution central moments for different division strategies *λ* ∈ {1, 2, 0.5} and division steps *M* ∈ {1, 10}. **A**., **B**. and **C**. Mean cell size ⟨*s*⟩ as a function of time *t*. *s* is measured in units of expected size at birth 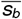 and *t* is measured in terms of the doubling time *τ*. **D**.,**E**. and **F**. Fluctuations in cell size *CV*^2^(*s*) as a function of time. Solid line: Numerical solution of (10) through integration of (4). The shaded region corresponds to the 95% confidence interval of the simulations of five thousand cells. (Parameters: *µ* = ln(2), *s*_0_ = 1, *CV*^2^(*s*_0_) = 0. *k*_*d*_ is selected as 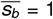. For *λ* = 1, *k*_*d*_ = *M* ln(2); for *λ* = 0.5, *k*_*d*_ ≈ 0.84*M*; and for *λ* = 2, *k*_*d*_ ≈ 0.46*M*).

Figure 3 shows the predictions obtained by solving (4) numerically. The results of Monte Carlo simulations (SSA) in five thousand cells are also presented. Note that oscillations appear when additional division steps *M* > 1 are considered. These oscillations are robust and do not decay over time. As discussed in previous articles, these oscillations appear as a result of the symmetry properties of the system [9, 10, 31].

Currently, there is no way to accurately measure how many cycle stages *m* a cell has completed. To overcome this issue, as presented in Figure 4, we can take advantage of the independence of the cycles and the robustness of growth and the rates of division in steady growth. In this way, we can manually synchronize all the trajectories so that the cells begin their dynamics approximately just after some arbitrary division (Figure 4B and Figure 4C). This technique has previously been used by other researchers [20, 28]. Thus, we can start with the assumption of *m*(*t* = 0) = 0 such as *P*_*m,n*_(*t* = 0) = *δ*_*m*,0_*δ*_*n*,0_ and compare the experimental trajectories with those observed in Figure 3.

**Figure 4:**
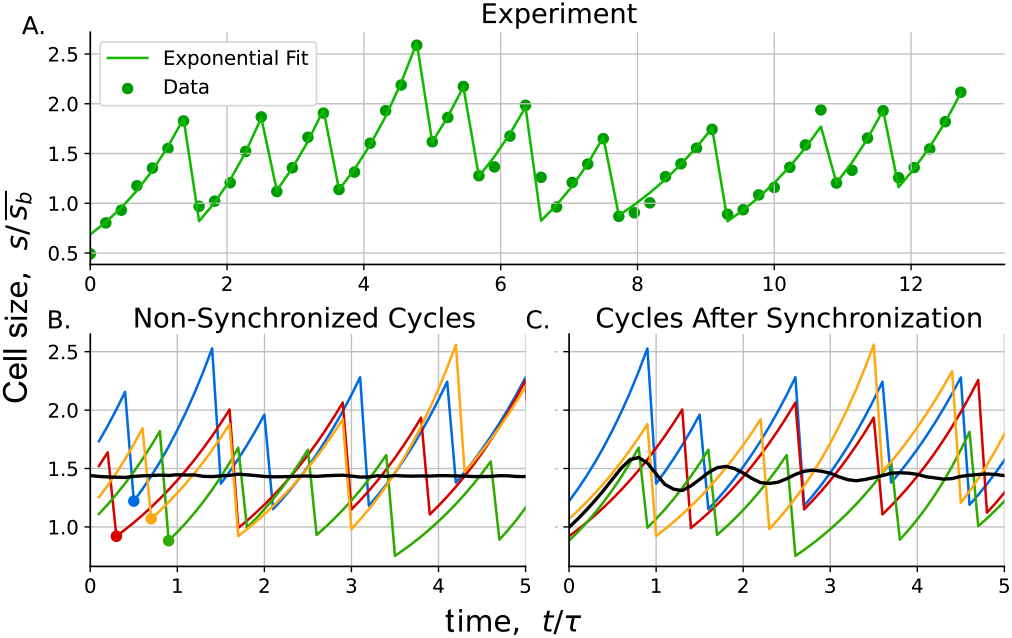
Single cell size trajectories as function of time. **A**. Example of an experimental trajectory (dots) and exponential fit (solid line). **B**. Multiple trajectories in which the size after the most recent division is highlighted with a point. **C**. These size trajectories are synchronized so that they start from their most recent division. After synchronization, the mean size, represented by the black line, now exhibits oscillations.

## Additional sources of noise

Until now, we have shown how to estimate the cell-size distribution considering stochasticity coming only from the division timing. Figure 3 shows how the main prediction is that moments show robust oscillations over time. In this section, we will show how to add other sources of noise and observe how they affect the dynamics of the moments of the cell size distribution.

### The heterogeneity of starting size

In general, cells do not start their cycles with the same size *s*_0_ in (7). Let *ρ*(*s*_0_) be the PDF of the cell size at *t* = 0; thus, the PDF *ρ*(*s*) at an arbitrary time follows:

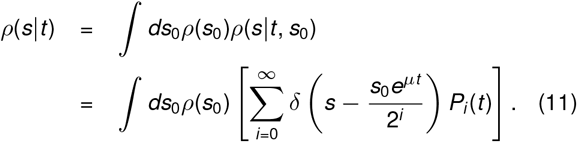

Figures 5A and 5D show how the oscillation amplitude of the central moments of the cell size is lower than the amplitude obtained considering a specific size (7). This occurs because the terms in the sum (11) are out phase. If the initial noise *CV*^2^(*s*_0_) is high enough, the oscillations disappear and the system evolves into a steady-moment distribution. The noise level, that is, the noise around which the dynamics oscillates, does not change. This means that *CV*^2^(*s*_0_) does not contribute to *CV*^2^(*s*).

**Figure 5:**
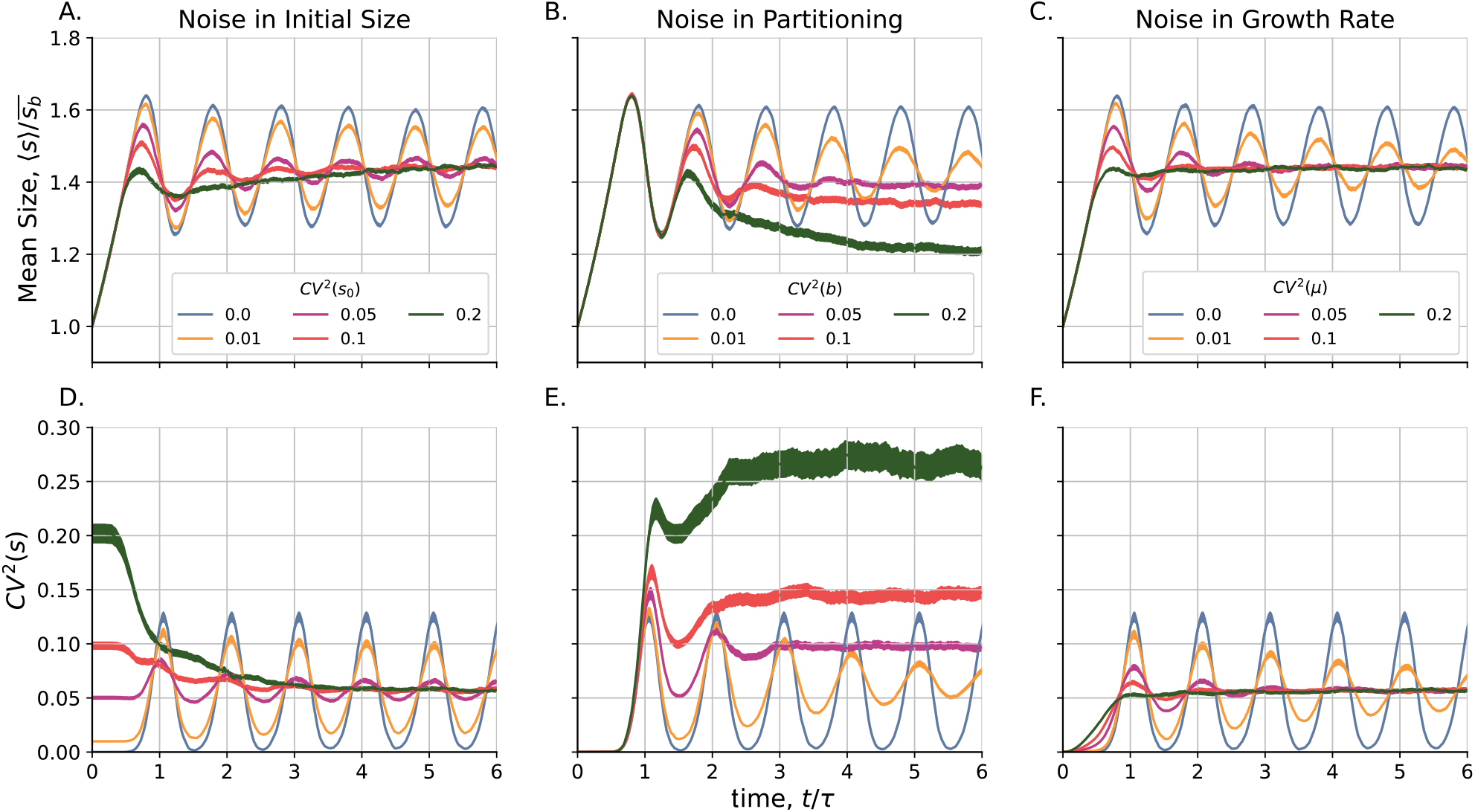
Effects of some sources of noise in the size distribution moment dynamics. **A**.,**B**. and **C**. Mean size as a function of time. Cell size is measured in units of 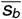, the mean size at birth. **D**., **E**. and **F**. Cell size fluctuations measured by the square coefficient of variation *CV*^2^(*s*) = var(*s*)*/*⟨*s*⟩^2^. The line width represents the 95% confidence interval of simulations (Parameters: 10000 cells, *M* = 20, *µ* = ln(2), *k*_*d*_ = 20*ln*(2). To study THE noise in the initial size, we consider initial sizes as gamma-distributed variables with mean ⟨*s*_0_⟩ = 1 and specified *CV*^2^(*s*_0_). To study noise in partitioning position, we consider the partitioning ratio *b*, the ratio between the size of daughters and mothers cells, as a beta-distributed variable with mean ⟨*b*⟩ = 1*/*2 and specified *CV* (*b*). To study the noise in the growth rate, we consider the growth rate *µ* as a constant per cycle and an i.i.d. variable selected from a gamma distribution with mean ⟨*µ*⟩ = ln(2) and specified *CV*^2^(*µ*). The corresponding division rate *k*_*d*_ is also selected for each cycle, such as 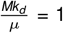)

### Noise in partitioning position

If we consider that the cells are not exactly divided by half as assumed in (2). As explained in Figure 2A, we can implement simulations by, after each division, multiplying the size times a random variable *b* with ⟨*b*⟩ = 0.5 and variability *CV*^2^(*b*) > 0. Using a process similar to (2), we can conclude that the size after *n* divisions at time *t* follows [9]

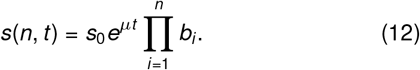

Phenomenologically, *b* can be approximated to a beta-distributed variable with variance measured from experiments [23].

Figures 5B and 5 E show the mean size ⟨*s*⟩ and the cell size variability *CV*^2^(*s*) over time considering noise in the splitting position *CV*^2^(*b*). With increasing *CV*^2^(*b*), the oscillation becomes more damped and the steady mean size decreases. As discussed in [9], the steady mean size can decrease, since the noisier the splitting, the more likely it is to have small cells. The cycles of these small cells last longer, given the exponential nature of growth. Therefore, these cells are overrepresented in a random population. Regarding fluctuations in cell size, the increase in *CV*^2^(*b*) also increases *CV*^2^(*s*), as suggested by other authors [23].

### Cell-to-cell noise in growth rate

Assume that cells do not grow at the same rate *µ* as explained in (1). Instead, as explained in Figure 2D, suppose that after a division *j*, a cell grows during that cycle with rate *µ*_*j*_, being an i.i.d. variable from a distribution centered on *µ* and noise *CV*^2^(*µ*). We neglect the correlation in growth rate for successive cycles [20, 23, 32] or the correlation with other variables such as size at birth [32]. Numerically, we modeled these *µ* as independent gamma-distributed variables centered on the mean growth rate ⟨*µ*⟩, using the variance observed experimentally. Returning to the noiseless splitting regime, the size *s* at time *t* after *n* divisions, each occurring at instant *t*_*j*_ is now:

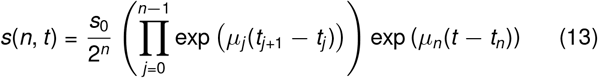

Figures 5C and 5F show the effect of increasing *CV*^2^(*µ*) while keeping the ratio *k*_*j*_/*µ*_*j*_ constant. That is, once the division rate *k* cannot be measured *a priori*, it is selected to be proportional to the growth rate. As a main result, we observed that the central moments of the size distribution exhibit damped oscillations, but *CV*^2^(*µ*) does not affect the main level of *CV*^2^(*s*). This observation was already observed in recent studies [15].

### Cell size autocorrelation Function (ACF)

The autocorrelation function (ACF) tells us about the relationship between the cell size at a time *t* and its own size after a time *t* + *t′*. To estimate the ACF, we first make sure that the size distribution moments are steady and do not oscillate over time. For simulations, we collect data from cells starting from a noisy size distribution *CV*^2^(*s*_0_) ≈ 0.2 after they reach stationary moments. Experimentally, as can be seen in Figure 4B, if the size trajectories are not synchronized, the distribution moments do not show oscillations (see also Figure S1).

Setting *t* = 0 at 2 h after imaging acquisition (see Figure S1) and knowing the size *s*_*i*_ (*t*) of the cell *i* at a time *t*, ACF *γ*(*t′*) is estimated using the formula:

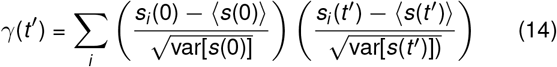

In the models, neither ⟨*s*⟩ nor 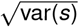 depend on time. However, as we shall discuss later, in the experiment, these moments do. However, the ratio var(*s*)*/*⟨*s*⟩2 is robust, justifying the use of (14).

## Experiments

To observe how oscillations in cell size moments behave in nature, we performed experiments on a *Mother Machine* microfluidic device (Figure 1C). We tracked the size dynamics of *E. coli* cells in minimal medium with two different carbon sources: glucose (2179 cells), a medium in which cells exhibit a strategy *adder* (*λ* = 1) and glycerol (665 cells), where the division strategy is *sizer-like* (*λ* = 1.5) [18, 33]. Single cell trajectories were synchronized with their most recent division, as explained in Figures 4B and 4C Details of the experiment can be found in Supplementary Information.

Once *λ* is adjusted from the division strategy graph ⟨Δ⟩ versus *s*_*b*_, the number of division steps can be estimated from the added size noise [18]. The division rate *k*_*d*_ can be estimated from the mean growth rate *µ* and the formula (6) or, if a general division strategy is considered, we fit the mean size at birth ⟨*s*_*b*_⟩ with 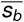. Variability in *s*_0_ was measured experimentally.

We assume no correlation between these random variables except the growth rate *µ* and the division rate *k*_*d*_. Although we try to avoid adjudicating on a specific biochemical mechanism, the most probable interpretation of *k*_*d*_ is the rate of synthesis of the FtsZ polymer [8]. On the other hand, *µ* is precisely the mean rate of protein synthesis. If we consider the absence of correlation between *µ* and *k*_*d*_, the mean size dynamics appears to be strongly damped compared to the oscillations observed in the experiments and shown in Figure 6. For simplicity, we assume that *µ* and *k*_*d*_ are proportional.

**Figure 6:**
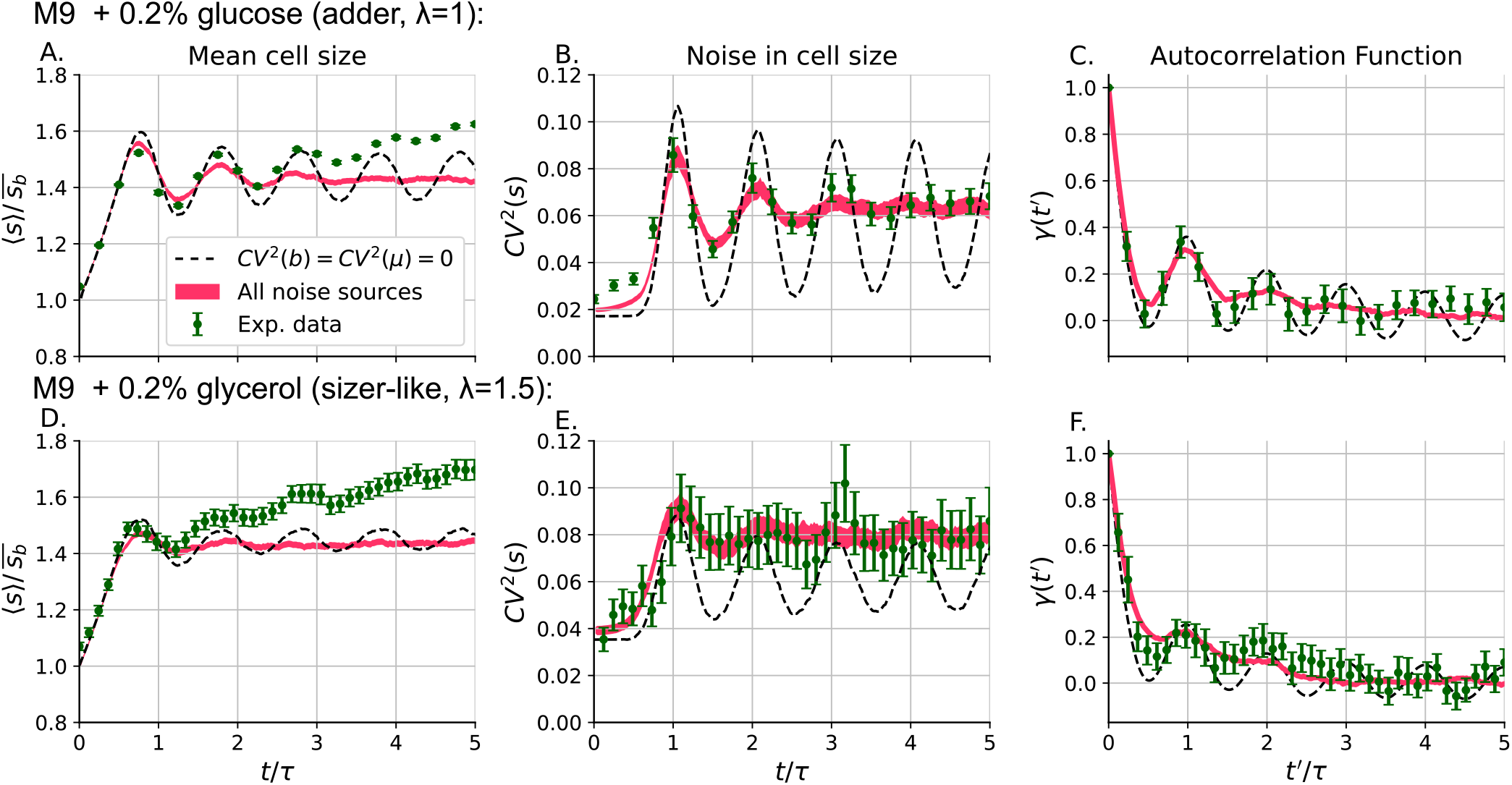
Robustness of the oscillations in the moments of the size distribution. **A**. Mean cell size, **B**. Cell size fluctuations, **C**. Cell size autocorrelation function (ACF), as functions of time for *E. coli* bacteria growing in minimal media with glucose as carbon source. **D. E**. and **F**. same as A., B. and C. but for cells growing in minimum medium with glycerol as the carbon source. The experimental results are shown by green dots with error bars representing the 95 % confidence intervals. The black dashed lines show the results of numerical solutions that used only noise in the cycle timing and initial size. The pink lines are the result of simulations with parameters fitted to the data. The width of the pink line also represents the 95% confidence interval of the simulations. 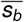 is the average cell size at birth. The parameters used to fit the experiments are shown in supplementary information.

Figure 6 shows the comparison between the dynamics of the predicted moments considering only noise in timing and initial size (black dashed line) and the experimental findings (green dots). In other words, we set *CV*^2^(*µ*) = *CV*^2^(*b*) = 0. Although we found oscillations in the central moments of the experimental cell size histogram, these oscillations are damped. To adjust these trends, we will next include additional sources of noise.

We selected cycles with an excellent fit to exponential growth (*R*^2^ > 0.9) to estimate *CV*^2^(*µ*) and *CV*^2^(*b*). Supplementary information (Table S2) presents the trends of these variables. After using the measured values in our model, we observed that the resulting oscillations should be more damped than the observed ones. We will discuss the origin of this discrepancy later. To overcome this issue, we fitted other parameters. The *CV*^2^(*b*) was estimated from the level of *CV*^2^(*s*) as explained in Figure 5E and *CV*^2^(*µ*) from the damping of the oscillations in *CV*^2^(*s*) as observed in Figure 5F.

After the carbon source was changed to glycerol, we observed a higher level of noise. In our theory, this implies more damped oscillations and an increase in *CV*^2^(*s*) to a value higher than that predicted with simple noise (4), as shown in Figure 6E.

Our measurements suggest that although the growth rate did not change significantly during the experiment (see Figure S1B in SI), the mean cell size increased approximately 10% during the entire time lapse. A possible cause might be that the time after cell insertion was not long enough for the cells to finish adapting.

In Figure 6C (glucose) and Figure 6F (glycerol), we present the comparison between the experimental ACF *γ*(*t′*) obtained using (14) from simulations. We observe that the ACF in the experimental data is slightly higher than the one predicted by the simulations, but shows a similar damped oscillatory pattern. The difference may be due to the correlations between variables for successive cycles (not considered in the simulations) and the mean cell size increasing over time.

## Discussion

Cell size regulation models based on division rates are not well understood due to limited mappings to experimental results of cell size dynamics [9]. Here, considering that division is an stochastic process that occurs at a rate proportional to a power of the size, we explore its implications for division control. As noted previously [18, 27] this model can reproduce most known division strategies, that is, the relation between the added size (Δ = *s*_*d*_ − *s*_*b*_) and the size at birth (*s*_*b*_).

We modeled the dynamics of the size distribution of a population of independent cells. After the cell size trajectories are synchronized starting from their most recent division, our model predicts oscillations at the central moments of the cell size and the cell size autocorrelation function (ACF). These oscillations are robust even considering stochasticity in the timing of division and the initial cell size. Experimentally, although the oscillations are evident, they are damped. From our theory, we found this damping after considering additional noise sources (noise in both division position and growth rate).

If we do not consider any correlation between the division rate constant *k*_*d*_ in (3) and the growth rate *µ* in (1), we observe that the oscillations should be strongly damped. Since the damping was slight, we assume that *k*_*d*_ and *µ* are proportional. A possible justification is that *k*_*d*_ is related to the rate of accumulation of a molecule (such as FtsZ) to trigger division [8] and that there is generally a relationship between protein synthesis and growth rate [34, 35].

Another intriguing effect shown in Figures 6A and 6D is that the average cell size increases despite the fact that the average growth rate remains constant. This is probably because the cells did not recover fully from insertion into the microfluid. We can obtain a better fit to the data if it is assumed that *k*_*d*_ changes over time (see Figure S3 in SI). The non-constant *k*_*d*_ is surprising because it implies that cells can reach a steady growth rate (*µ*) before stability in division *k*_*d*_. Additional research on these nonstationary processes, for instance, the size variation in changing environments such as the growth curve [36, 37] or other kind of controlled growth conditions [38], can give us more details about the relationship between the variables of the size dynamics.

We observe how size control is altered after increasing the level of noise in biochemical processes inside the cell as a response to stress. In our case, this stress can increase when bacteria are allowed to grow in a poor nutrient medium (M9 + glycerol) [39]. In this medium, there is an increase in noise both in growth rate and in partitioning position. As an effect of this noise increasing, we predict that the oscillations in the cell size moments should be more damped. Additional experiments can further study our hypothesis. Using mutant bacteria with greater variability in the position of the septum, for example, bacteria with a mutation in the Min system [40], can quantify the effect of noise in the split position without changing other biochemical noise. Another option may be to adjust the noise in gene expression by adapting the bacteria to a minimal medium [41, 42]. For example, a strain adapted to growth medium in glycerol as a carbon source could exhibit less noise in gene expression and thus less damped oscillations.

Although we observe this desynchronization in the cell cycles, the observed damping is less than expected from the measured noises in both the splitting position and the growth rate. This suggests that there are additional hidden mechanisms to control the timing of the cycle. Some of them can include correlations between growth rate and size at birth (see supplementary information), as recently reported [32], mechanisms associated with chromosome replication to trigger division [43, 44], or other hidden variables relative to environmental conditions or intrinsic factors of each particular lineage [45]. We expect to face this more complex approach in future research.

Some researchers have shown that gene expression noise can be helpful in cells as a source of variability in phenotypes [46–48], and that the resulting diversified strategies (bet hedging) can be advantageous from an evolutionary perspective [49, 50]. Our results hint at a subtler but related advantage of intracellular fluctuations. By causing rapid desynchronization of the cell cycles of sister cells, they can maintain population-level homogeneity in metabolic rates, which could help maintain homeostasis. The impact of this size regulation on cell proliferation will be part of the study in future articles. Some additional variables must be incorporated into the model, such as the correlation between cell lineages [51]. Therefore, it is not yet clear whether this result will become an evolutionary advantage.

Determining the underlying mechanisms in division control allows us to understand not only cell growth, but also signaling within cells, which depends on concentrations [52]. These concentrations can fluctuate depending on the timing of cell division and its variability [23,53]. Therefore, having an accurate stochastic model of cell division is essential to predict phenotypic variability and control intracellular circuits. Specifically, some of these applications included recently proposed frameworks for gene expression analysis [4,54] and cell lineage analysis of experimental data from proliferating cell populations [55, 56].

## Supplementary Information (SI)

### SI Datasets

The data set and the script used for the data analysis can be found online [57] (https://doi.org/10.5281/zenodo.691769) and (https://github.com/canietoa/SizeDynamics).

### SI Movies

We present the dynamics of cell distribution in the supplementary videos1 and video2.

### Strain

*A*ll strains used in this study are *Escherichia coli* k-12 MG1655 background [58].

### Plasmid construction

We obtained a plasmid with GFP-mut2 under the promoters pNac or pRpoD from the Uri Alon Plasmid Library [59] and we modified the plasmid to insert a constitutive RFP promoter, induced by RNA1, as a segmentation marker. As a backbone, we used the pUA66 plasmid from the Uri Alon library [59] and linearized it with a BglII restriction enzyme -Thermo scientific. The insert, the constitutive marker mCherryKate2, was then amplified from the DHL60 strain from the Paulsson Lab at Harvard Medical School using colony PCR. The final assembly was performed using the Gibson assembly protocol of New England BioLabs.

### Growth media

Defined medium was used in all experiments. For *Escherichia coli*, we used M9 minimal medium [60] with different carbon sources, glucose or glycerol, as shown in Table S1.

**Table S1:**
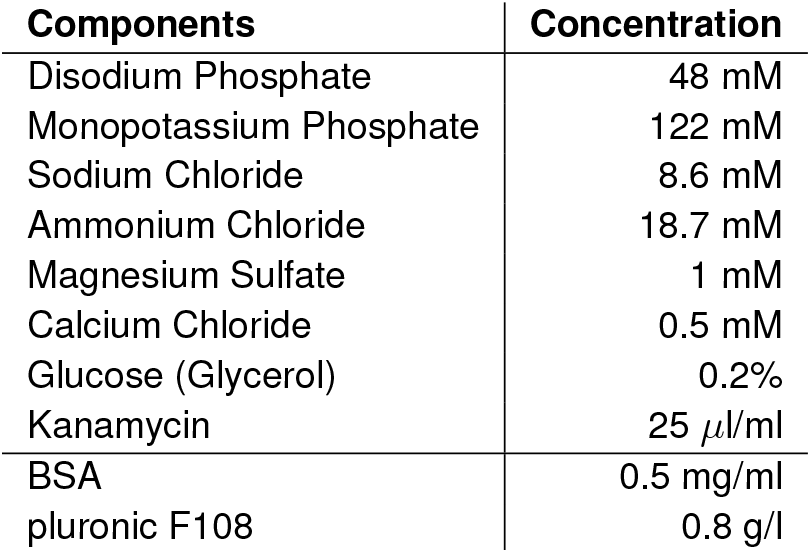
M9 medium details.

### Cell preparation

Before each time-lapse image, cells were selected from a single colony on an agar plate that was streaked no more than 7 days prior to use. Cells were inoculated in M9 with selection antibiotics; in our case, kanamycin 25 ul / ml. After 12-18 hours at 37 ^*o*^ C on a water bath shaker, cells were diluted 1,000-fold into 2 mL of the same defined medium as that used in the microfluidic experiment. After shaking at 37^*o*^C in a water bath until OD600 = 0.1-0.4, cells were diluted again 100- to 1,000-fold into the same medium and shaken at 37 ^*o*^C in a water bath until OD600 = 0.2. The cell culture was then concentrated 10- to 100-fold and injected into a microfluidic device *Mother Machine* via a micropipette with gel loading tips. Furthermore, 0.5 mg / ml BSA -Bovine serum albumin, Gemini Bio Products, CA-was added to the fresh growth medium to reduce cell adhesion to the surface of microfluidic channels. The medium was then added to 60 ml plastic syringes -BD- and flowed using a syringe pump with a flow of 30 *µl/min* for time-lapse imaging. All imaging experiments were conducted at 37 ^*o*^C in an environmental chamber.

### Microfluidics

*Mother Machine* microfluidic devices were used to monitor single cell growth for 12-20 generations. Master molds, from each of which many PDMS microfluidic devices were cast, were manufactured using standard nanofabrication techniques (detailed protocols are available in “The Mother Machine Handbook” from the Jun lab website at http://jun.ucsd.edu and a video at http://www.youtube.com/watch?v=RGfb9XU5Oow), courtesy of the Paulsson lab. PDMS was prepared from a Sylgard 184 Silicone Elastomer kit: polymer base and curing agent were mixed in a ratio of 10 to 1, air bubbles were purged in a vacuum chamber, the degassed mixture was poured over the master, and the devices were cured for about 1h at 90^*o*^C. Cured PDMS has a rubber-like consistency that allows devices to be peeled manually from the master mold. Devices were treated with Isopropyl Alcohol to remove residual uncured polymer from the PDMS matrix.

**Figure S1:**
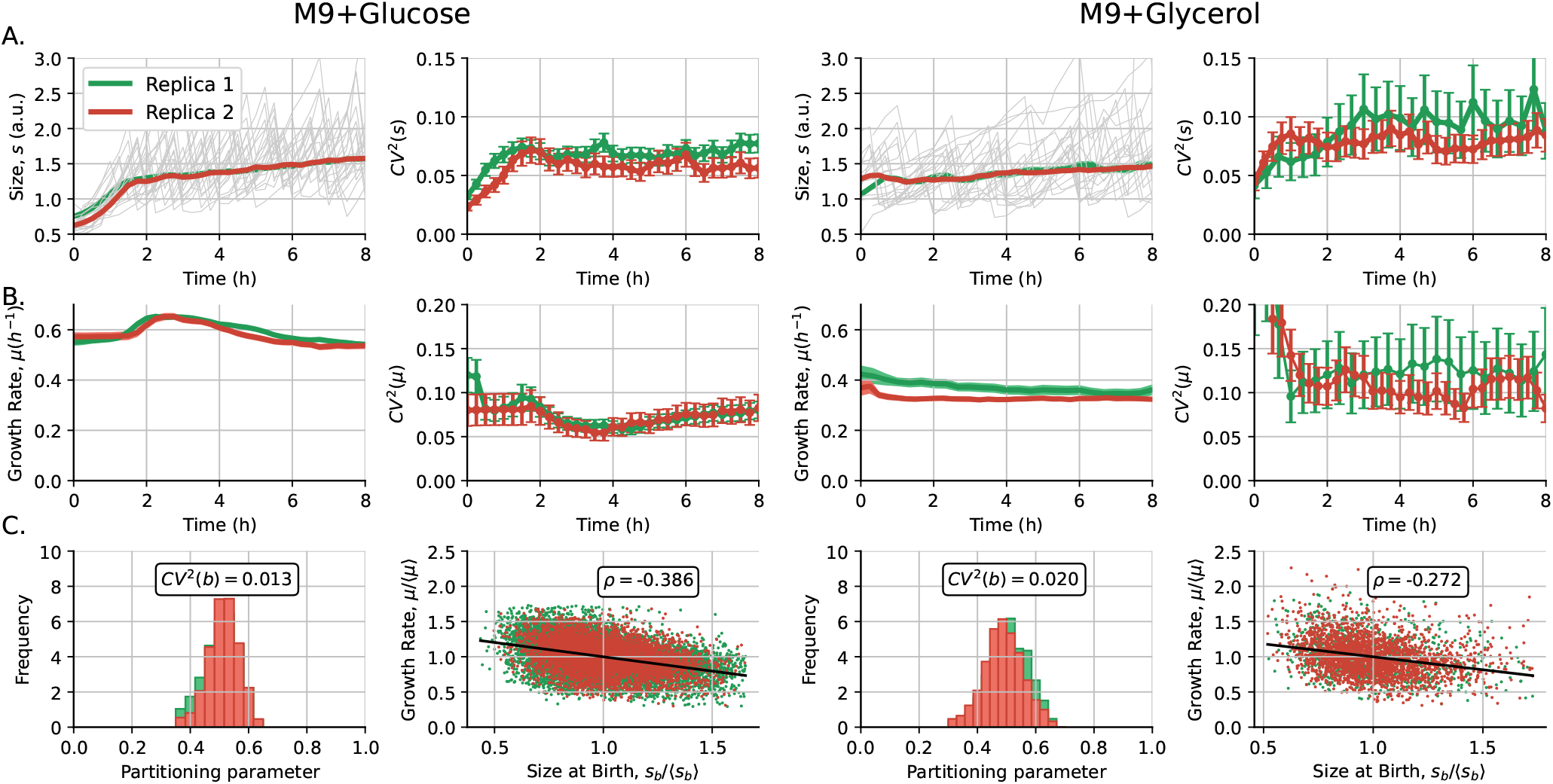
Measured variables of size regulation. **A**. Cell size and size variability *CV*^2^(*s*) as a function of time for M9 + glucose (left) and M9 + glycerol (right). **B**. Mean growth rate and its variability *CV*^2^(*µ*) as a function of time for M9 + glucose (left) and M9 + glucose (right). **C**. Histogram of the partitioning ratio *b* and the relationship between growth rate *µ* and size at birth *s*_*b*_ for M9 + glucose (left) and M9 + glycerol (right). Different colors represent the two replicas taken for each growth condition.

To bond the PDMS layers, the surfaces were exposed to oxygen plasma for 15 seconds at 30 watts in a Harrick Plasma system. Oxygen plasma makes exposed PDMS and glass reactive, so that covalent bonds form between surfaces brought into contact with one another. The seal between the PDMS surfaces was established for 10 minutes at 65 ^*o*^C.

### Microscopy and image acquisition

An inverted Nikon Eclipse Ti microscope equipped with a Perfect Focus system, a 60x air objective lens (NA 0.95), a lumencor spectra X3 light engine and an Andor Zyla 4.2 PLUS sCMOS camera were used for fluorescense imaging. The filter set used was the ET-Sedat Quad-band (8900, Chroma Technology Crop). The exposure time was set to 200 ms and the illumination intensity was set at 100%. The time-lapse frequency was 15 minutes or 22 minutes.

### Image analysis

Briefly, the segmentation was done using images from a bright, constitutively expressed RFP on a PRNA1 promoter. The rough trench boundaries were estimated with the Otsu threshold method followed by erosion, opening, and dilation of the mask. Then the binding box of the trenches found was used to find the cells within. The cells were then segmented in each of the trenches using Niblack segmentation [61]. Cells joined by their poles (as indicated by objects with definite restrictions) were separated using the top 10% brightest pixels of cells as a seed for the watershed. Spurious noncell objects were rejected using their size, orientation, and shape. Finally, the boundaries were refined using opening, thickening, and active contours. The parameters chosen for each experiment required extensive testing, and segmentation was manually checked. We chose to follow only the cells at the closed end of the channel.

### Fitting the size dynamics to simulations

To compare the experimental trajectories of the central moments of the size distribution with simulations, we normalize the experimental doubling time to that we can theoretically consider the doubling time *τ* = 1. This condition sets the mean growth rate ⟨*µ*⟩ = ln(2), which is also normalized. The division strategy was considered to fit the parameter *λ* in the SRF (3) *h* = *k*_*d*_ *s*^*λ*^ to the experimental trend of Δ vs. *s*_*b*_. The size was normalized by the experimental mean size at birth, which is associated with theoretical 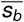 which, with the constraint 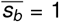 and given the division strategy, the mean value of the constant ⟨*k*_*d*_⟩ can be inferred using numerical rootfinders. To simplify the model, we consider that the particular *k*_*d*_ in a given cell cycle was proportional to the growth rate (considered stochastic). Noise in the initial size 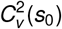 was measured. The other resulting fitting parameters are stored in Table S1.

The low value of *M* for bacteria that grow in Glycerol compared to the *M* obtained from bacteria that grow in glucose can be due to additional sources of noise such as decorrelation not measured between the growth rate *µ* and the division rate *k*_*d*_ that, for simplicity, were assumed to be proportional in each cycle. The division steps can also change if the actual division mechanism explaining the sizer-like strategy is different from the power law.

### Measured parameters of cell regulation

Figure S1 shows the measured variables of cell regulation. Figure S1 shows the size dynamics and size variability as a function of the time before cycle sinchronization for both growth conditions. Figure S1B shows the dynamics of the mean growth rate and the fluctuations around this value over time. These trajectories help us estimate the noise in growth rate *CV*^2^(*µ*) as shown in Table A2. Figure S1C also presents the histogram of the partitioning ratio *b* for the cycles studied, and their fluctuations over the mean *CV*^2^(*b*) are also presented. The correlation between growth rate *µ* and size at birth *s*_*b*_ was a parameter that was not considered in the modeling in the main article. We measured this correlation and presented it in Figure S1C.

**Table S1:**
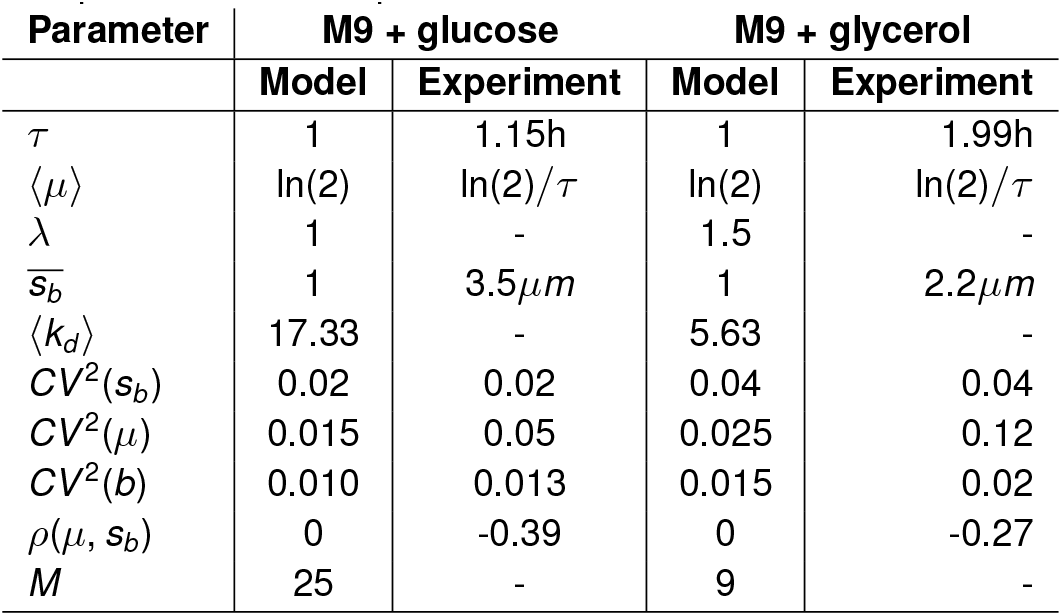
Fitting parameters adjusted to the trajectories of the central moments of the size distribution. Measured noises are also presented as comparison.

| Parameter | M9 + glucose |  | M9 + glycerol |  |
| --- | --- | --- | --- | --- |
|  | Model | Experiment | Model | Experiment |
| $\tau$ | 1 | 1.15h | 1 | 1.99h |
| $\langle \mu \rangle$ | $\ln(2)$ | $\ln(2)/\tau$ | $\ln(2)$ | $\ln(2)/\tau$ |
| $\lambda$ | 1 | - | 1.5 | - |
| $\overline{s_b}$ | 1 | $3.5\mu m$ | 1 | $2.2\mu m$ |
| $\langle k_d \rangle$ | 17.33 | - | 5.63 | - |
| $CV^2(s_b)$ | 0.02 | 0.02 | 0.04 | 0.04 |
| $CV^2(\mu)$ | 0.015 | 0.05 | 0.025 | 0.12 |
| $CV^2(b)$ | 0.010 | 0.013 | 0.015 | 0.02 |
| $\rho(\mu, s_b)$ | 0 | -0.39 | 0 | -0.27 |
| $M$ | 25 | - | 9 | - |

**Figure S2:**
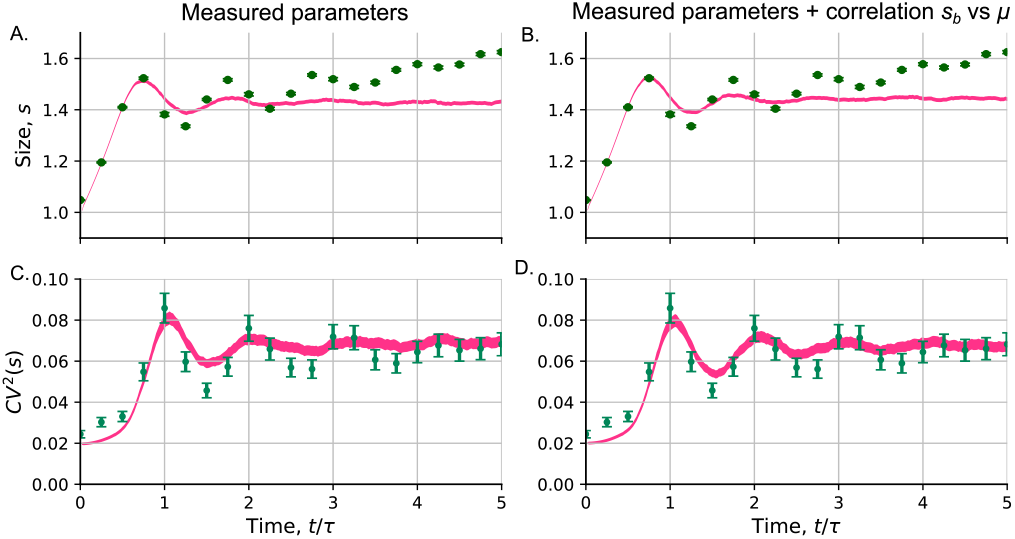
Effects of adding correlations between size regulation variables in the fitting of the moments dynamics. **A**. Mean size and **C**. size fluctuations *CV*^2^(*s*) over time considering the measured noise. **B**. Mean size and **D**. size fluctuations *CV*^2^(*s*) over time considering measured noise, including also the correlation between growth rate *µ* and size at the beginning of the cell cycle *s*_*b*_. Comparison between experiments of cells that grow in M9 with glucose as the carbon source (green dots) and simulations (pink line).

### The effects of hidden correlations

Figure S2A shows that, considering the measured parameters of cell regulation (Table S1), the expected oscillations in the size moments seem to be more damped than the observed ones.

We believe that a complex model can improve the fit of the model and the data. As recently reported [32], there are some correlations between cell regulation variables that are not yet very well studied. In our case, as an example, we study the effects of the correlation between growth rate and size at birth S2B. We can see that, after including this correction, the expected oscillations in the moments are less damped.

### The effects of a variable division rate

**Figure S3:**
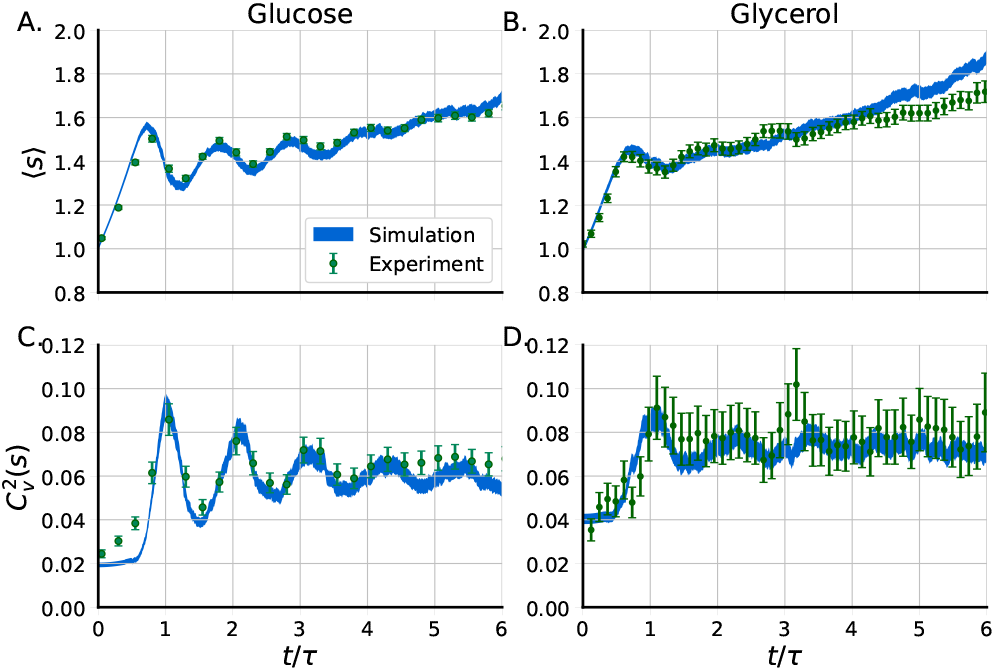
Fitting of the experimental size dynamics considering division rate *k*_*d*_ variable over time. **A**. Mean size and **C**. size fluctuations *CV*^2^(*s*) for cells growing in M9 with glucose as the carbon source. **B**. Mean size and **D**. size fluctuations *CV*^2^(*s*) over time for cells growing in M9 minimal medium with glycerol as the carbon source. Comparison between experiments of cells that grow in M9 with glucose as the carbon source (green dots) and simulations (blue line).

To overcome the issue of an increase in the mean size over time, we propose that the division rate *k*_*d*_ was not constant during the time lapse. Figure S3 shows how, after proposing that *k*_*d*_ increases linearly with time, keeping the growth rate constant, the fit to the data is improved.

## Acknowledgements

We thank the Paulsson lab at Harvard Medical School for making their laboratory available for some of our measurements and image analysis. The authors also thank COLCIENCIAS convocatoria para doctorados nacionales 647 and for financial support.

## Author contributions statement

C.N and C.V. Perform theoretical details and simulations. A.S., C.V. and C.N. Conceive the theory. J.C.A.C conducted the experiment, C.S. performed the image analysis, C.N. analyzed the data, and J.P. designed the experiments and directed the research. All authors reviewed the manuscript.

## References

[1] Y.-M. Zhang and C. O. Rock, “Membrane lipid homeostasis in bacteria,” Nature Reviews Microbiology, vol. 6, no. 3, pp. 222–233, 2008.

[2] N. Demaurex, “ph homeostasis of cellular organelles,” Physiology, vol. 17, no. 1, pp. 1–5, 2002.

[3] H. Kempe, A. Schwabe, F. Crémazy, P. J. Verschure, and F. J. Bruggeman, “The volumes and transcript counts of single cells reveal concentration homeostasis and capture biological noise,” Molecular biology of the cell, vol. 26, no. 4, pp. 797–804, 2015.

[4] J. Lin and A. Amir, “Homeostasis of protein and mrna concentrations in growing cells,” Nature communications, vol. 9, no. 1, pp. 1–11, 2018.

[5] C. A. Vargas-Garcia, K. R. Ghusinga, and A. Singh, “Cell size control and gene expression homeostasis in single-cells,” Current opinion in systems biology, vol. 8, pp. 109–116, 2018.

[6] P. A. Levin and S. Taheri-Araghi, “One is nothing without the other: theoretical and empirical analysis of cell growth and cell cycle progression,” Journal of molecular biology, vol. 431, no. 11, pp. 2061–2067, 2019.

[7] S. Jun, F. Si, R. Pugatch, and M. Scott, “Fundamental principles in bacterial physiology—history, recent progress, and the future with focus on cell size control: a review,” Reports on Progress in Physics, vol. 81, no. 5, p. 056601, 2018.

[8] F. Si, G. Le Treut, J. T. Sauls, S. Vadia, P. A. Levin, and S. Jun, “Mechanistic origin of cell-size control and homeostasis in bacteria,” Current Biology, vol. 29, no. 11, pp. 1760–1770, 2019.

[9] C. Nieto, C. Vargas-Garcia, and J. M. Pedraza, “Continuous rate modeling of bacterial stochastic size dynamics,” Physical Review E, vol. 104, no. 4, p. 044415, 2021.

[10] P. Bokes and A. Singh, “Cell volume distributions in exponentially growing populations,” in International Conference on Computational Methods in Systems Biology, pp. 140–154, Springer, 2019.

[11] D. Huh and J. Paulsson, “Random partitioning of molecules at cell division,” Proceedings of the National Academy of Sciences, vol. 108, no. 36, pp. 15004–15009, 2011.

[12] M. Soltani, C. A. Vargas-Garcia, D. Antunes, and A. Singh, “Intercellular variability in protein levels from stochastic expression and noisy cell cycle processes,” PLoS computational biology, vol. 12, no. 8, p. e1004972, 2016.

[13] N. Nordholt, J. Van Heerden, R. Kort, and F. J. Bruggeman, “Effects of growth rate and promoter activity on single-cell protein expression,” Scientific reports, vol. 7, no. 1, p. 6299, 2017.

[14] D. J. Kiviet, P. Nghe, N. Walker, S. Boulineau, V. Sunderlikova, and S. J. Tans, “Stochasticity of metabolism and growth at the singlecell level,” Nature, vol. 514, no. 7522, p. 376, 2014.

[15] C. Jia, A. Singh, and R. Grima, “Cell size distribution of lineage data: analytic results and parameter inference,” Iscience, vol. 24, no. 3, p. 102220, 2021.

[16] P.-Y. Ho, J. Lin, and A. Amir, “Modeling cell size regulation: From single-cell-level statistics to molecular mechanisms and population-level effects,” Annual review of biophysics, vol. 47, pp. 251–271, 2018.

[17] S. Taheri-Araghi, S. Bradde, J. T. Sauls, N. S. Hill, P. A. Levin, J. Paulsson, M. Vergassola, and S. Jun, “Cell-size control and homeostasis in bacteria,” Current Biology, vol. 25, no. 3, pp. 385–391, 2015.

[18] C. Nieto, J. Arias-Castro, C. Sánchez, C. Vargas-García, and J. M. Pedraza, “Unification of cell division control strategies through continuous rate models,” Physical Review E, vol. 101, no. 2, p. 022401, 2020.

[19] M. Osella, E. Nugent, and M. C. Lagomarsino, “Concerted control of escherichia coli cell division,” Proceedings of the National Academy of Sciences, vol. 111, no. 9, pp. 3431–3435, 2014.

[20] P. Wang, L. Robert, J. Pelletier, W. L. Dang, F. Taddei, A. Wright, and S. Jun, “Robust growth of escherichia coli,” Current biology, vol. 20, no. 12, pp. 1099–1103, 2010.

[21] J. T. Sauls, D. Li, and S. Jun, “Adder and a coarse-grained approach to cell size homeostasis in bacteria,” Current opinion in cell biology, vol. 38, pp. 38–44, 2016.

[22] M. Priestman, P. Thomas, B. D. Robertson, and V. Shahrezaei, “Mycobacteria modify their cell size control under sub-optimal carbon sources,” Frontiers in cell and developmental biology, vol. 5, p. 64, 2017.

[23] S. Modi, C. A. Vargas-Garcia, K. R. Ghusinga, and A. Singh, “Analysis of noise mechanisms in cell-size control,” Biophysical journal, vol. 112, no. 11, pp. 2408–2418, 2017.

[24] M. Campos, I. V. Surovtsev, S. Kato, A. Paintdakhi, B. Beltran, S. E. Ebmeier, and C. Jacobs-Wagner, “A constant size extension drives bacterial cell size homeostasis,” Cell, vol. 159, no. 6, pp. 1433–1446, 2014.

[25] C. A. Nieto-Acuna, C. A. Vargas-Garcia, A. Singh, and J. M. Pedraza, “Efficient computation of stochastic cell-size transient dynamics,” BMC bioinformatics, vol. 20, no. 23, pp. 1–6, 2019.

[26] K. R. Ghusinga, C. A. Vargas-Garcia, and A. Singh, “A mechanistic stochastic framework for regulating bacterial cell division,” Scientific reports, vol. 6, p. 30229, 2016.

[27] C. A. Vargas-García and A. Singh, “Elucidating cell size control mechanisms with stochastic hybrid systems,” in 2018 IEEE Conference on Decision and Control (CDC), pp. 4366–4371, IEEE, 2018.

[28] M. Wallden, D. Fange, E. G. Lundius, Ö. Baltekin, and J. Elf, “The synchronization of replication and division cycles in individual e. coli cells,” Cell, vol. 166, no. 3, pp. 729–739, 2016.

[29] S. Iyer-Biswas, C. S. Wright, J. T. Henry, K. Lo, S. Burov, Y. Lin, G. E. Crooks, S. Crosson, A. R. Dinner, and N. F. Scherer, “Scaling laws governing stochastic growth and division of single bacterial cells,” Proceedings of the National Academy of Sciences, vol. 111, no. 45, pp. 15912–15917, 2014.

[30] G. Micali, J. Grilli, M. Osella, and M. C. Lagomarsino, “Concurrent processes set e. coli cell division,” Science advances, vol. 4, no. 11, p. eaau3324, 2018.

[31] E. Bernard, M. Doumic, and P. Gabriel, “Cyclic asymptotic behaviour of a population reproducing by fission into two equal parts,” arXiv preprint arXiv:1609.03846, 2016.

[32] M. Kohram, H. Vashistha, S. Leibler, B. Xue, and H. Salman, “Bacterial growth control mechanisms inferred from multivariate statistical analysis of single-cell measurements,” Current Biology, vol. 31, no. 5, pp. 955–964, 2021.

[33] C. Nieto, J. Arias-Castro, C. Vargas-Garcia, C. Sanchez, and J. M. Pedraza, “Noise signature in added size suggests bacteria target a commitment size to enable division,” bioRxiv, 2020.

[34] S. Klumpp, Z. Zhang, and T. Hwa, “Growth rate-dependent global effects on gene expression in bacteria,” Cell, vol. 139, no. 7, pp. 1366–1375, 2009.

[35] M. Scott, C. W. Gunderson, E. M. Mateescu, Z. Zhang, and T. Hwa, “Interdependence of cell growth and gene expression: origins and consequences,” Science, vol. 330, no. 6007, pp. 1099–1102, 2010.

[36] S. Bakshi, E. Leoncini, C. Baker, S. J. Cañas-Duarte, B. Okumus, and J. Paulsson, “Tracking bacterial lineages in complex and dynamic environments with applications for growth control and persistence,” Nature Microbiology, vol. 6, no. 6, pp. 783–791, 2021.

[37] T. Shimaya, R. Okura, Y. Wakamoto, and K. A. Takeuchi, “Scale invariance of cell size fluctuations in starving bacteria,” Communications Physics, vol. 4, no. 1, pp. 1–12, 2021.

[38] F. Büke, J. Grilli, M. C. Lagomarsino, G. Bokinsky, and S. J. Tans, “ppgpp is a bacterial cell size regulator,” Current Biology, vol. 32, no. 4, pp. 870–877, 2022.

[39] J. Marles-Wright and R. J. Lewis, “Stress responses of bacteria,” Current opinion in structural biology, vol. 17, no. 6, pp. 755–760, 2007.

[40] V. W. Rowlett and W. Margolin, “The bacterial min system,” Current biology, vol. 23, no. 13, pp. R553–R556, 2013.

[41] M. Richard and G. Yvert, “How does evolution tune biological noise?,” Frontiers in genetics, vol. 5, p. 374, 2014.

[42] J. Liu, J.-M. François, and J.-P. Capp, “Use of noise in gene expression as an experimental parameter to test phenotypic effects,” Yeast, vol. 33, no. 6, pp. 209–216, 2016.

[43] G. Witz, E. van Nimwegen, and T. Julou, “Initiation of chromosome replication controls both division and replication cycles in e. coli through a double-adder mechanism,” Elife, vol. 8, p. e48063, 2019.

[44] M. Berger and P. R. t. Wolde, “Replication initiation in e. coli is regulated via an origin-density sensor generating adder correlations,” arXiv preprint 2106.03674, 2021.

[45] H. Vashistha, M. Kohram, and H. Salman, “Non-genetic inheritance restraint of cell-to-cell variation,” Elife, vol. 10, p. e64779, 2021.

[46] J. M. Raser and E. K. O’shea, “Noise in gene expression: origins, consequences, and control,” Science, vol. 309, no. 5743, pp. 2010–2013, 2005.

[47] M. B. Elowitz, A. J. Levine, E. D. Siggia, and P. S. Swain, “Stochastic gene expression in a single cell,” Science, vol. 297, no. 5584, pp. 1183–1186, 2002.

[48] H. Maamar, A. Raj, and D. Dubnau, “Noise in gene expression determines cell fate in bacillus subtilis,” Science, vol. 317, no. 5837, pp. 526–529, 2007.

[49] K. Lewis, “Persister cells,” Annual review of microbiology, vol. 64, pp. 357–372, 2010.

[50] J.-W. Veening, W. K. Smits, and O. P. Kuipers, “Bistability, epigenetics, and bet-hedging in bacteria,” Annu. Rev. Microbiol., vol. 62, pp. 193–210, 2008.

[51] M. ElGamel, H. Vashistha, H. Salman, and A. Mugler, “Multigenerational memory in bacterial size control,” arXiv preprint 2206.05340, 2022.

[52] A. Camilli and B. L. Bassler, “Bacterial small-molecule signaling pathways,” Science, vol. 311, no. 5764, pp. 1113–1116, 2006.

[53] Z. Vahdat, Z. Xu, and A. Singh, “Modeling protein concentrations in cycling cells using stochastic hybrid systems,” IFAC-PapersOnLine, vol. 54, no. 9, pp. 521–526, 2021.

[54] J. Jakub Jedrak, M. Kwiatkowski, and A. Ochab-Marcinek, “Exactly solvable model of gene expression in a proliferating bacterial cell population with stochastic protein bursts and protein partitioning,” Phys. Rev. E, vol. 99, p. 042416, Apr 2019.

[55] R. García-García, A. Genthon, and D. Lacoste, “Linking lineage and population observables in biological branching processes,” Phys. Rev. E, vol. 99, p. 042413, Apr 2019.

[56] P. Thomas and V. Shahrezaei, “Coordination of gene expression noise with cell size: analytical results for agent-based models of growing cell populations,” Journal of the Royal Society Interface, vol. 18, no. 178, p. 20210274, 2021.

[57] C. Nieto, “Sizedynamics,” July 2022.

[58] K. F. Jensen, “The escherichia coli k-12” wild types” w3110 and mg1655 have an rph frameshift mutation that leads to pyrimidine starvation due to low pyre expression levels.,” Journal of bacteriology, vol. 175, no. 11, pp. 3401–3407, 1993.

[59] A. Zaslaver, A. Bren, M. Ronen, S. Itzkovitz, I. Kikoin, S. Shavit, W. Liebermeister, M. G. Surette, and U. Alon, “A comprehensive library of fluorescent transcriptional reporters for escherichia coli,” Nature methods, vol. 3, no. 8, p. 623, 2006.

[60] J. H. Miller, Experiments in molecular biology. 1972.

[61] O. Samorodova and A. Samorodov, “Fast implementation of the niblack binarization algorithm for microscope image segmentation,” Pattern Recognition and Image Analysis, vol. 26, no. 3, pp. 548–551, 2016.

